# clonealign: statistical integration of independent single-cell RNA & DNA-seq from human cancers

**DOI:** 10.1101/344309

**Authors:** Kieran R Campbell, Adi Steif, Emma Laks, Hans Zahn, Daniel Lai, Andrew McPherson, Hossein Farahani, Farhia Kabeer, Ciara O’Flanagan, Justina Biele, Jazmine Brimhall, Beixi Wang, Pascale Walters, IMAXT Consortium, Alexandre Bouchard-Côté, Samuel Aparicio, Sohrab P Shah

## Abstract

Measuring gene expression of genomically defined tumour clones at single cell resolution would associate functional consequences to somatic alterations, as a prelude to elucidating pathways driving cell population growth, resistance and relapse. In the absence of scalable methods to simultaneously assay DNA and RNA from the same single cell, independent sampling of cell populations for parallel measurement of single cell DNA and single cell RNA must be computationally mapped for genome-transcriptome association. Here we present `clonealign`, a robust statistical framework to assign gene expression states to cancer clones using single-cell RNA-seq and DNA-seq independently sampled from an heterogeneous cancer cell population. We apply `clonealign` to triple-negative breast cancer patient derived xenografts and high-grade serous ovarian cancer cell lines and discover clone-specific dysregulated biological pathways not visible using either DNA-Seq or RNA-Seq alone.

---

Recent advances in genomic measurement technologies have allowed for unprecedented scalable interrogation of the genomes and transcriptomes of single-cells (Zahn et al., 2017; Zheng et al., 2017). Such technologies are of particular interest in cancer, enabling measurement of cell-autonomous properties which constitute tumours as a whole. Molecular phenotyping at the single-cell level enables reconstruction of tumour life histories through phylogenetic analysis (Jahn et al., 2016; Smith et al., 2017), assessment of cell types in the tumour microenvironment (Schelker et al., 2017), and quantification of intra-tumoural heterogeneity and its clinical implications (Tellez-Gabriel et al., 2016; Mitra et al., 2014).

Theoretically, combined assays sequencing both RNA and DNA from the same single cell will provide a measurement of genomic alterations impacting transcriptional programmes. This would yield a powerful single-cell level genotype-phenotype read out, encoding relevant malignant properties of clonal expansion, proliferation and metastasis. Moreover, drug responses in cancer are commonly driven by positive and negative evolutionary selection of mutation-induced phenotypes, but genome-independent responses via dynamic epigenetic re-wiring of transcriptional programs have also been observed (Shaffer et al., 2017). Thus, multimodal approaches assaying both DNA and RNA are essential for comprehensive study of drug response. While pioneering technologies such as G&T-seq (Macaulay et al., 2015) and DR-seq (Dey et al., 2015) sequence both the DNA and RNA from single-cells, they measure few cells compared to assays that sequence DNA or RNA alone such as Direct Library Preparation (DLP, Zahn et al. (2017)) or 10X genomics single-cell RNA-seq (Zheng et al., 2017), and thus provide only a limited view of each tumour’s genomic and transcriptional heterogeneity.

Independent analysis however, introduces a new analytical challenge in how to associate cells from independently measured experiments. Assuming a population structure with a fixed number of clones, this can be expressed as a mapping problem, whereby cells measured with transcriptome assays must be aligned to those measured with a genome assay. To address this problem we introduce `clonealign`, a statistical method to assign cells measured with single-cell RNA-seq to clones derived from low-coverage single-cell DNA-seq (figure 1A). In our approach, clones are defined at the genomic level by building a phylogenetic tree using single-cell copy number measurements. In order to relate the independent measurements (figure 1B), we assume that an increase in the copy number of a gene will result in a corresponding increase in that gene’s expression and vice versa (figure 1C), a relationship previously observed in joint RNA-DNA assays in bulk tissues (Curtis et al., 2012) and at the single-cell level (Macaulay et al., 2015; Dey et al., 2015; Han et al., 2018). Based on this relationship we formulate a statistical model that explains the observed gene expression pattern in terms of the copy number profile of a clone present in the scDNA-seq data and thus assigns each cell to a clone (methods).

**Figure 1:**
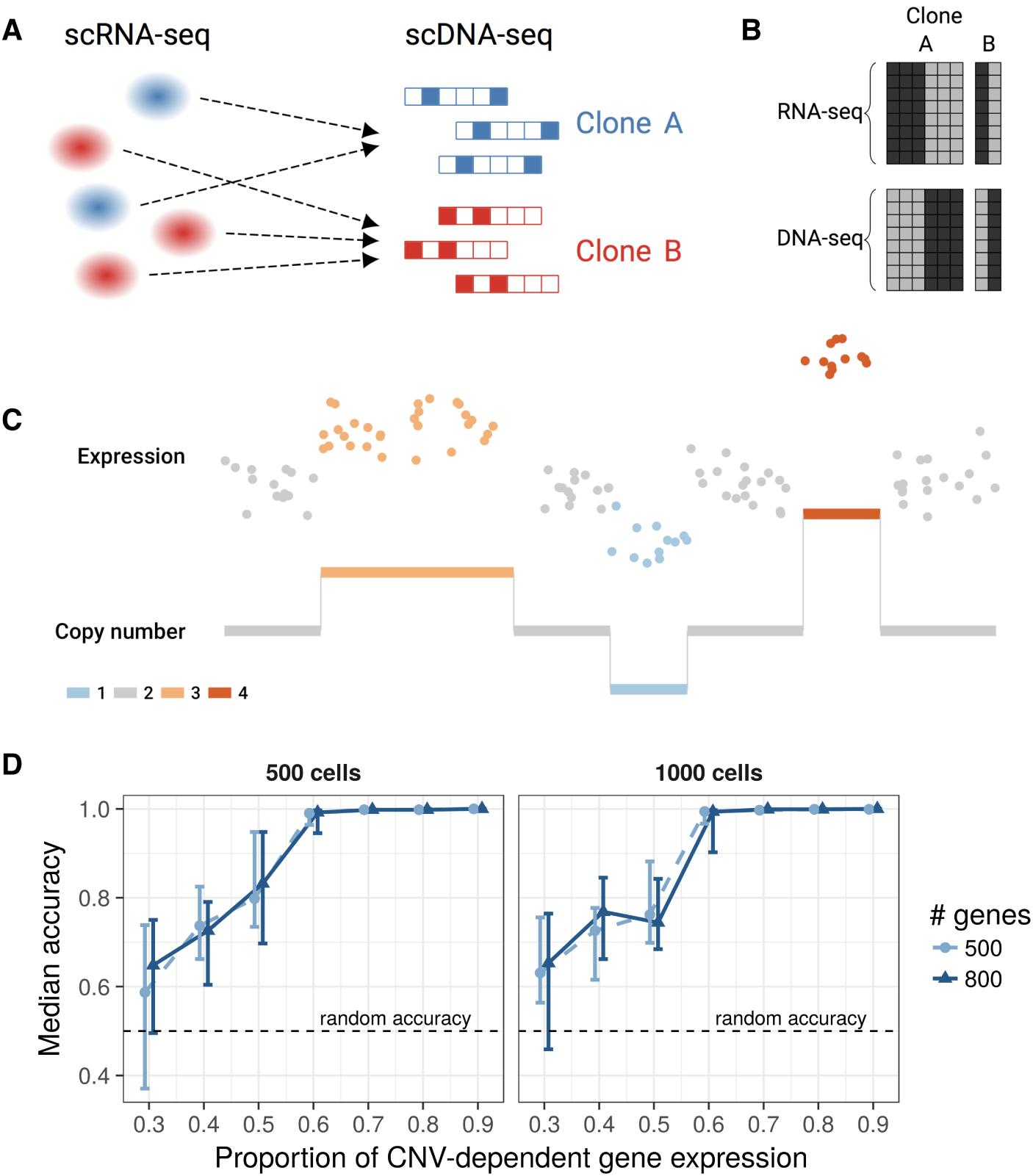
Assigning single-cell RNA-seq to clone-of-origin using clonealign. **A** Given independently sampled single-cell DNA- and RNA-seq from the same tumour, we would to assign each cell in RNA-seq space to its clone-of-origin in DNA-space and thus infer clone-specific gene expression. **B** The current scalable measurement technologies do not allow for simultaneous measurement of RNA and DNA in the same cell, meaning although the same clones will be present we will never encounter the same cell in different sequencing assays. **C** To relate cells as measured in RNA-space to their clones measured in DNA-space we assume a relationship between gene copy number and gene expression. **D** Comprehensive simulations demonstrate the robustness of `clonealign` to the underlying proportion of genes exhibiting a copy number dosage effect. If more than 40% of gene expression has a copy number dependence clone assignment is significantly better than random, while at more than 50% clone assignment is close to perfect.

To test the robustness the `clonealign` model, we performed comprehensive simulations (methods) where a certain proportion of genes were simulated assuming no CN-expression relationship and clone assignments re-inferred using `clonealign` assuming all genes had CN-dependent expression. We found that `clonealign` is highly robust to variation in the underlying proportion of genes with CN-dependent expression (figure 1D and supplementary figure 1), even when up to 50% of genes do not exhibit CN-dependent expression.

We next investigated the capacity of our approach to reveal clone-specific phenotypic properties in real cancer data, using the serially passaged triple-negative breast cancer patient derived xenograft SA501 as a substrate. SA501 exhibits a complex clonal architecture and reproducible clonal dynamics over successive xenograft passages (Eirew et al., 2015). Thus it is an ideal model system to exemplify clone-specific gene expression. We previously described single-cell DLP DNA-seq for SA501X3F (Zahn et al., 2017), a copy number analysis of which identified three genotypically-distinct clones (A, B, & C) with prevalences 82.3%, 10.8%, and 6.9% respectively, with clone A further expanding in subsequent passages.

We linked gene expression to clones in SA501 by generating single-cell RNA-seq from the SA501X2B xenograft passage using 10X genomics (methods) and assigned each cell to a clone (A, B or C) using `clonealign`. 1152 single-cells post-QC (methods) were assigned to clones A, B, and C with prevalence of 80.6%, 13.8%, and 5.6%, closely matching the expected proportions inferred from the single-cell DNA-seq (82.3%,10.8%, and 6.9%). A genome-wide view of the clone-specific copy number and expression profiles reveals a strong dosage effect as modelled by `clonealign` in all but one region (figures 2A&B). The clone assignments are highly confident for clone A but some cells exhibit uncertainty of assignment between clones B & C (figure 2C), reflecting a combination of having more cells in clone A as well as more similar expression profiles of B & C but distinct expression profiles of (B or C) relative to A. This latter explanation is further supported in a t-SNE analysis (Maaten and Hinton, 2008) using only genes residing in chromosome regions with variable copy number between clones (figure 2D).

**Figure 2:**
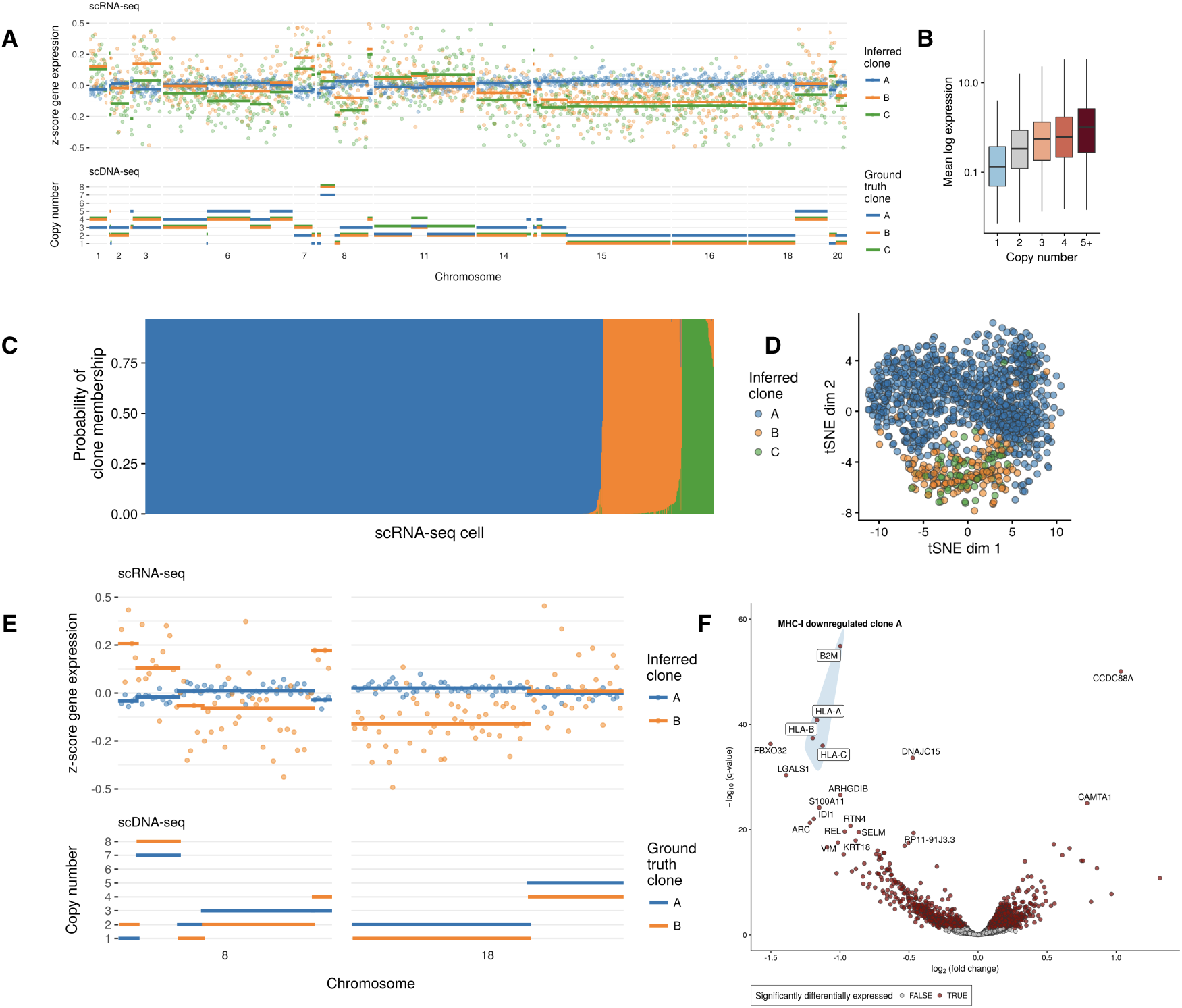
Inferring clone-dependent gene expression in SA501 triple-negative breast cancer xenograft. **A** Clone-specific copy number for ground truth clones in scDNA-seq (bottom) and clone-specific *z*-score expression for `clonealign` inferred clones in scRNA-seq (top) for regions exhibiting inter-clone copy number aberrations. In every copy number segment except one, when the copy number for a given clone is higher than others then on average the normalized gene expression is also higher. **B** The mean log expression as a function of copy number across all clones. **C** Clone assignment probabilities for 1152 single-cell RNA-seq profiles across three clones. `clonealign` confidently assigns cells to clone A, with some cells exhibiting high assignment uncertainty between clones B & C. **D** A tSNE analysis using only genes residing in copy number regions shows the cells clustering by clone. **E** *z*-score normalized gene expression and copy number profiles for held-out data on chromosomes 8 & 18 as a function of genomic position (gene index along chromosome). In all but one copy number segment, when the copy number profile of a clone is higher the normalized gene expression in that chromosome is also higher on average. **F** Differential expression analysis for genes residing in regions whose copy number is identical between clones highlights downregulation of MHC class I proteins.

We next sought to validate the `clonealign` assignments by both testing the internal consistency of our model and with a held out, orthogonal data source. We re-inferred the clones for SA501X2B using genes from all chromosomes except 8 and 18. If both the clone assignments and the expression-CNA assumption are correct then the expression of genes on the held-out chromosomes (8 & 18) should closely correlate with the copy number profiles of those chromosomes. In all-but-one copy number segments of the held-out chromosomes, congruency between copy number levels and normalised gene expression was observed: where the copy number profile of a clone was higher, the normalized gene expression in that chromosome was also higher and vice-versa (figure 2E). We formulated this into a statistical test asking if given the clone assignments and copy number profiles we can predict the expression of genes on the held-out chromosomes better than can be expected at random, with a null distribution established over permuted clone assignments. Comparing `clonealign` clone assignments to the null distribution with RMSE of predictions showed significantly better predictive accuracy than could be expected at random (*p* < 10^−3^, supplementary figure 2). We then added a further validation measure using a loss-of-heterozygosity (LOH) analysis (methods) to discover if clone-specific LOH events observed in DNA space were also observed in RNA space. A single allele resulting from a genomic LOH event can only yield mono-allelically expressed transcripts (Ha et al., 2012). Although the allele frequency data were sparse and low coverage at germline heterozygous sites, we observed an LOH event on chromosome 18 in clone B which was mono-allelically expressed in the scRNA-seq (supplementary figure 3). Finally we quantified the robustness of `clonealign` to input gene selection by incrementally reducing the number of input genes and comparing the consistency in clone assignment to the assignments using all genes (methods), finding up to 60% of genes could be retained and the resulting clone assignments remain highly consistent (supplementary figure 4).

Having established the validity of the clone assignments, we next sought to determine clone-specific phenotypes using gene expression as a proxy. We performed a differential expression analysis comparing cells assigned to clone A to those assigned to clones B & C using Limma voom (Law et al., 2014). 47% of genes (510/1084) residing in clone-specific copy numbers (CSCN) regions were differentially expressed compared to 23% of genes in regions with identical copy number (ICN) regions (1905/8201). Clone A is distinguished by loss of an entire X-chromosome, but it was previously unknown whether the loss constituted the active or inactive copy. We observed downregulation of X-inactive specific transcript *X I ST* (supplementary figure 5) - expressed only on the inactive X chromosome - in clone A, implying the retained chromosome is the active copy.

We next examined the differential expression of genes residing in regions with identical copy number between clones. By construction these genes would not be impacted by gene dosage *in cis*, but may be altered through signalling networks *in trans* where upstream transcriptional regulators lie in copy number altered regions. We found systematic downregulation of the MHC class I cell surface proteins in clone A (figure 2F and supplementary figure 6) along with *β*_2_ microglobulin (*B2M*), suggesting a clone-specific deficiency in presenting intra-cellular proteins to cytotoxic T cells, and therefore a putative mechanism by which clone A progressively dominates the SA501 xenograft tumours in subsequent passages. Loss of MHC expression is a mechanism of tumour immune escape (Garrido et al., 2016, 2012) and our results indicate this may be selected for despite the immune-deficient environment of the murine host. Importantly, clone A did not exhibit LOH in any HLA gene in clone A (supplementary figure 7), implying MHC class-I downregulation is due to transcriptional pathway alterations.

We supplemented our differential expression analysis with a variance components analysis (Arnol et al. (2018) & methods) to partition gene expression variation into either clone-specific or intrinsic / residual. This revealed genes whose expression variation was governed by genomic state (clonality), such as CD44 antigen - a marker of tumorigenic cancer cells (Al-Hajj et al., 2003) - of which around a quarter of expression variation is clone-specific (supplementary figure 8). To elucidate which pathways show clone-dependent regulation, we performed a gene set enrichment analysis (Subramanian et al., 2005) on all genes ranked by proportion of regulation explained by genomic state. Clone-specific immune response (figure 2F), including pathways involved in MHC class I mediated antigen presentation were highly ranked. To discover if any transcriptional states existed within clone assignments, we performed an intra-clonal clustering of the scRNA-seq data using SC3 (Kiselev et al., 2017) with *k* = 2 clusters and called cell cycle states using Cyclone (Scialdone et al., 2015). We found clusters within each clone largely separated based on G2M score (supplementary figure 9), implying the largest source of intra-clonal variation corresponds to cell cycle stage.

We next applied `clonealign` to DLP scDNA-seq and 10X genomics scRNA-seq data from two clonally related high grade serous carcinoma (HGSC) cell lines, derived from both ascites (`OV2295R`) and solid tumour (`TOV2295R`) at relapse from the same patient (Létourneau et al., 2012). We constructed a single-cell phylogeny on the derived copy number profiles from DLP+ using a Latent Tree Model (Farahani, 2018), yielding four distinct clades (figure 3A). We assigned the cells as measured using scRNA-seq to the DLP+ clones using `clonealign`, and found 1356 (40.9%) mapping to `TOV2295R_A`, 1960 (59.1%) to `TOV2295R_B`, 490 (33.6%) to `OV2295R_C`, and 970 (66.4%) to `OV2295R_D` (figure 3B, top). To ensure the clone assignments were accurate we tested whether predicted clone-specific expression of genes on held out chromosome segments correlated well with the copy number profiles of those genes (figure 3C & supplementary figures 10-12), and found these assignments to be robust to the choice of input gene (supplementary figures 13-14). Differential expression analysis on OV2295R identified 267/1122 (23.8%) in CSCN regions and 893/4455 (20.0%) as differentially expressed in ICN regions (supplementary figure 15), while a similar analysis in TOV2295R identified 953/1504 (63.4%) and 2423/5720 (42.3%) in CSCN and ICN regions respectively (supplementary figure 16).

**Figure 3:**
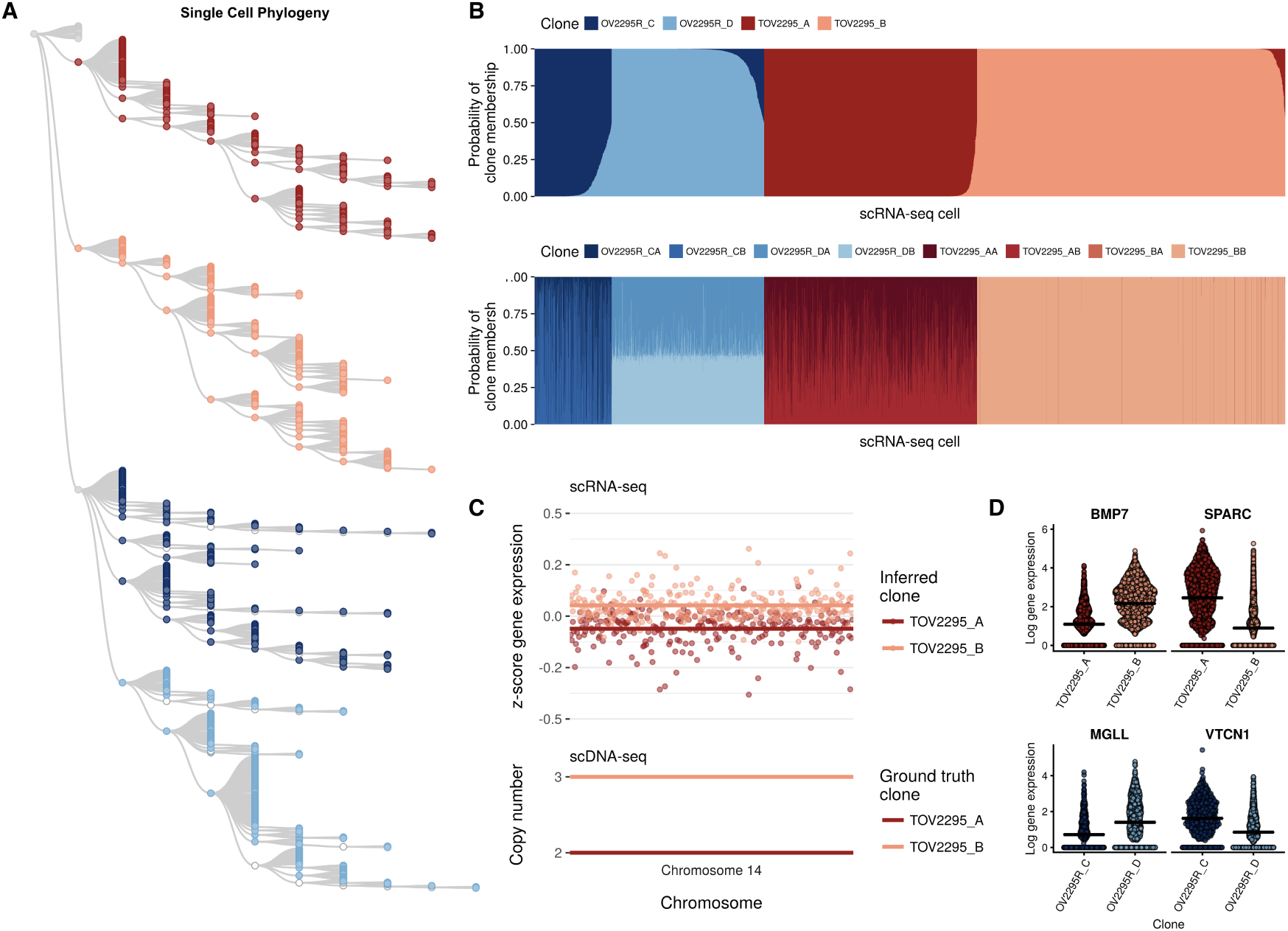
Clone-specific gene expression in high-grade serous ovarian cancer cell lines. **A** Single-cell phylogeny for the OV2295R and TOV2295R HGSC cell lines inferred using a Latent Tree Model divided into four clones (`TOV2295R_A, TOV2295R_B, OV2295R_C, OV2295R_D`). **B** The scRNA-seq clone assignments for the four clone model (top), then sub-divided into eight clones (bottom). **C** Expression-CNA relationship on two held out chromosomes for TOV2295R validates the `clonealign` fit. **D** Clone-dependent gene expression across OV2295R and TOV2295R

We next examined the ability of `clonealign` to resolve mappings as a function of phylogenetic distance between clones. In this analysis, higher levels of uncertainty in mappings between closely related clones is expected, assuming more closely related cells harbour more similar expression programs. Genomically defining a clone ultimately depends on clade-level groupings of cells that are approximately similar as a function of phylogenetic distance. We assembled a second set of clones from the OV2295R-TOV2295R phylogeny by sub-dividing each of the initial 4 clones into two (supplementary figure 17), and re-assigning each scRNA-seq cell to one of the 8 clones (figure 3B, bottom). We then computed Euclidean distance of each clone to its nearest neighbour and clone assignment probability for each cell. We found - as expected - a strong anti-correlation between the similarity of clones in genome space and the certainty with which cells are assigned to them (supplementary figure 18), demonstrating the analytical challenges of segregating cells into highly similar clones based on gene expression data alone. We further repeated the intra-clonal clustering analysis (as above for SA501), clustering each clone into two distinct groups separately and computing cell cycle phases. As with SA501, we found that in three of the four clones resultant clusters corresponded to cell cycle phase (supplementary figures 19 & 20), implying the largest genome-independent source of expression variation corresponds to cell cycle stage.

Our results establish a scalable statistical framework for assigning cells measured using scRNAseq to cancer clones measured independently using shallow scDNA-seq. We expect this approach can be used ubiquitously in the field of single cell biology including extensions for other multimodal approaches such as methylation-transcription and chromatin accessibility-transcription. At the edge of the field, sparse *in situ* measurements of transcription integrated with independent disaggregated sampling of single cell genomes are providing a route to studying spatial context of co-located cell populations. Finally, emergence of commercial platforms whereby single cell, kit-based assays for methylation, transcription and genome copy number are becoming widely available to the research community. In all of these settings, `clonealign` and future derivatives will provide a statistical framework to help interpret the cellular constituents of cancer, their fitness, and their phenotypes.

## METHODS

### CLONEALIGN: MODEL FORMULATION & INFERENCE

We begin with an *N × G* matrix of expression raw read counts **Y** for *N* cells and *G* genes, and a *G × C* matrix **Λ** = (*λ_gc_*) of clone specific copy numbers for *C* clones and *G* genes. Such a copy number matrix is typically obtained by phylogenetic analysis of single-cell CNV data, followed by cutting of the phylogenetic tree to produce *C* clones or clades. The goal of inference is to assign each of the *N* cells as measured in RNA-space to one of the *C* clones as measured in DNA-space.

For each cell *n* = 1, …, *N* we introduce a categorical assignment variable *z_n_* defined such that

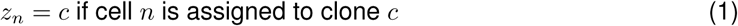

for *c* = 1, …, *C*. Our assumption is that *y_ng_* - the expression of gene *g* in cell *n* - will be dependent on the copy number of the gene in the clone to which *n* is assigned, ie 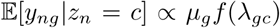 where *µ_g_* is the per-chromosome expression of gene *g* and *f* is a *dosage function* that maps the copy number of a gene to a multiplicative factor of expression. While this function is *a priori* unknown and joint estimation with clonal populations would lead to an unidentifiable model, we can encode some basic assumptions about gene dosage into our specification of *f*. We assume that if the copy number change is small it will lead to a proportional change in expression, e.g. a copy number of 3 could conceivably lead to 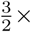 more expression. Conversely, we assume that if the copy number change is large, e.g. if a clone has copy number 12 in a particular region, the cells will have a compensatory mechanism such that fewer than 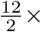 transcripts are produced, and that this is capped at an upper limit. With these considerations in mind we specify f as 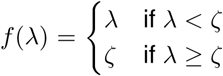, where in our analyses we fix *ζ* = 4. We leave as future work more sophisticated approaches such as inferring *f* from joint genomic-transcriptomic assays or marginalizing out *ζ* in Bayesian models.

We next consider specifying the exact likelihood model for `clonealign`. There is a subtlety in modelling RNA-seq data as outlined in Robinson and Oshlack (2010) in that the expression of each gene is measured relative to all other genes in a given library multiplied by the sequencing depth of that library. Taking this into account is of critical importance to our problem as if a highly expressed gene sits in a high copy number region in a clone it will cause a *decrease* in expression of all other genes. Therefore, the expected count of gene *g* in cell *n* conditional on that cell being assigned to clone *c* is given by

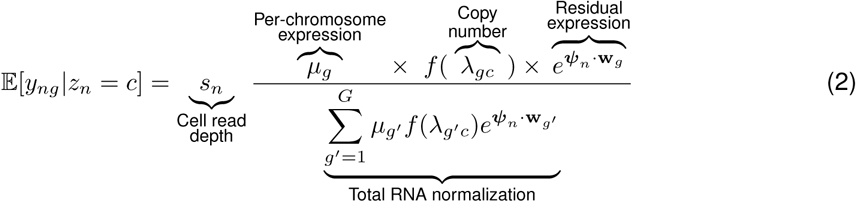

where *s_n_* is the total library size of cell *n*. We have introduced a *N* × *Q* matrix **Ψ** with row vectors *ψ_n_* and a *G* × *Q* matrix **W** with row vectors w*_g_* that we multiply together to produce a low-rank matrix of random-effects that helps explain residual gene expression variation in a similar manner to factor analysis. We set *Q* = 1 and ensure the model is weakly identifiable by imposing priors 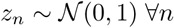.

We use a negative binomial likelihood as is commonly used to model both RNA-seq (Robinson and Oshlack, 2010; Love et al., 2014) and single-cell RNA-seq data (Risso et al., 2017). The mean of the negative binomial is given by Equation (2) with gene and clone specific dispersions *ϕ_cg_*, *g* = 1, …, *G*, *c* = 1, …, *C*. Estimating Negative Binomial dispersions can be unstable for low sample sizes which depends on clonal frequencies and is therefore unknown *a priori*. To increase the robustness of this estimation to the presence of rare clones we incorporate a hierarchical shrinkage prior 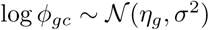 for *σ*^2^ > 0, where *η* is inferred from the data and *σ*^2^ is set by the user (we set *σ*^2^ = 1 for SA501 and TOV2295R while *σ*^2^ = 0.1 for OV2295R). The model as defined in 2 is invariant to rescalings of all *µ* so we fix *µ*_1_ = 1 and the interpretation of the remaining *µ*_2_, …, *µ_G_* is the per-chromosome expression relative to gene 1. The total library size *s_n_* can either be jointly inferred with the model or fixed beforehand. In all our analyses we set *s_n_* as the total counts of the genes considered multiplied by the TMM normalization (Robinson and Oshlack, 2010) factor calculated using the total library (across all measured genes).

We perform maximum a-posteriori inference of the model parameters **s**, ***µ***, ***ϕ***, **Ψ**, **W**, ***η*** as well as the clone assignment probabilities 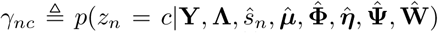 using an Expectation-Maximization algorithm (Dempster et al., 1977). Given *θ*^(*t*)^ denotes the value of parameter *θ* and EM iteration *t*, for each E-step we compute the clone assignment probabilities

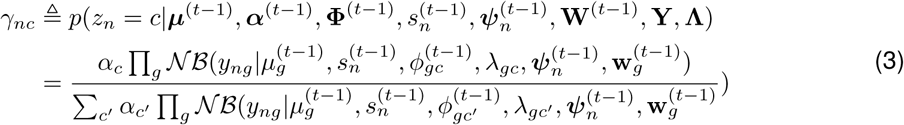

and form the *Q*-function

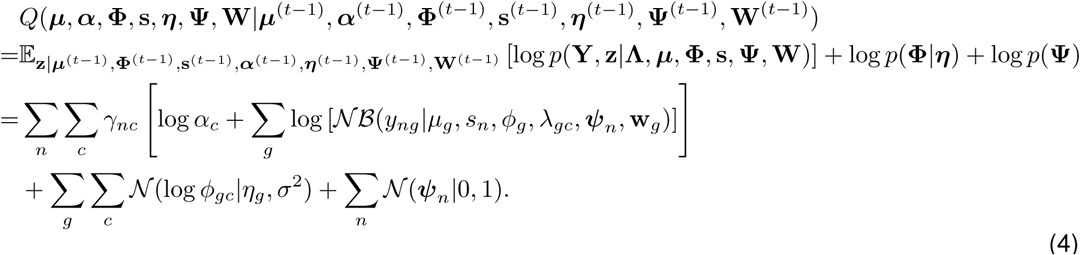

We typically set 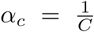 for both simulations and real data (where *α_c_* is the marginal probability that a cell originates from clone *c*), though such parameters may also be jointly inferred. The M-step involves the optimization of the *Q*-function with respect to the model parameters for which there is no analytic solution. Given the high dimensional nature of the optimization problem 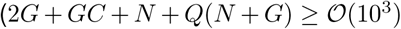 parameters) we turn to state-of-the-art high-dimensional optimization techniques designed for deep neural networks, using the Adam optimizer (Kingma and Ba, 2014) as implemented in Tensorflow. Convergence is assessed by monitoring the log marginal likelihood 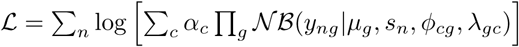. We found through emprical experimentation that model convergence is fastest using only a single ADAM update at each EM iteration, equivalent to performing gradient descent on the log marginal likelihood (Berg-Kirkpatrick et al., 2010). `clonealign` is open source and available online at http://www.github.com/kieranrcampbell/clonealign.

### SIMULATIONS

To ensure the simulations were as realistic as possible clonealign was fitted to the SA501 dataset giving an empirical distribution of the model parameters and data *p*(**Φ**, ***µ***, **Λ**)*p*(s). We then simulated from the clonealign model, sampling from the empirical distribution of model parameters for *N* = 500,1000 cells and *G* = 500,800 genes for genes with copy number ∈ 1, 2, 3, 4. For each simulation, a certain proportion *π* = 0.3, 0.4, …, 0.9 of genes were simulated with a CN-expression dependency, while the expression of the remaining 1− *π* proportion had an expression independent of copy number, achieved by setting the copy number to 2 for all clones during simulation of the expression, but providing the true copy number during inference as clonealign does not know *a priori* which genes exhibit a CN-expression dependency. Datasets were simulated for two clones corresponding to the A and B clones from SA501 with minor clone frequency uniformly simulated from [0.1, 0.5). Although clonealign infers clone-specific dispersion parameters these were set to be equal across clones in simulations (corresponding to the median dispersion inferred across clones from the joint density) to give a lower bound on model performance (though dispersions were allowed to be clone-specific during inference and *σ*^2^ was set to 1). During inference with clonealign all parameters were set to the defaults, with the metrics of accuracy (mean number of clone calls correct) as well as the area under the receiver-operator curve (AUC) being reported.

### BIOINFORMATICS ANALYSIS

For all scRNA-seq data expression estimates were obtained from raw read counts using CellRanger (version 2.0.1 for SA501X2B and version 2.1.0 for (T)OV2295R) aligned to hg19. Quality control of SA501X2B cells removed those with fewer than 1000 counts or 350 expressed genes in regions of distinct copy number between clones A, B & C. Clone specify copy number calls were created according to Zahn et al. (2017). X-chromosome genes were removed prior to clonealign analysis as the expression-copy number assumption will be violated if the deleted/amplified X copy is inactive. For OV2295R and TOV2295R cells were retained with total counts greater than 20000, and total number of genes detected between 3000 and 7500. Copy number calls for scDNA-seq were performed using HMMCopy version 1.22.0 and a phylogeny constructed using a latent tree model. The clone-specific copy number was constructed as the median copy number of all cells in a clone at a given position. Genes on the X-chromsome were removed as before.

Differential expression (DE) analysis was performed using Limma Voom (Law et al., 2014) version 3.36.0. For SA501X2B genes with greater than 100 total counts were retained for DE. For both OV2295R and TOV2295R genes with greater than 500 total counts were retained for DE as up to this threshold the mean-dispersion relationship reported by Limma Voom was visually a poor fit. All p-values were corrected for multiple hypothesis testing using the Benjamini-Hochberg procedure.

For the SA501 LOH analysis bulk whole-genome DNA sequencing as previously described in Eirew et al. (2015) was aligned to hg19 using BWA aln version 0.7.10 after which germline LOH alleles were identified using samtools 1.7 `mpileup` followed by VarScan 2.3.9 (Koboldt et al., 2012) `mpileup2snp` command (default settings). Single-cell RNA and DNA-seq profiles were merged into pseudobulk clones using samtools version 1.7 and reads mapping to ref and alt alleles at positions identified as germline heterozygous called using Varscan `mpileup2cns` command with default settings other than setting `-min-avg-qual 5` on the merged scRNA-seq to increase the number of callable positions. Regions in the pseudobulk pileups were called LOH using Titan version 1.16.0 (Ha et al., 2014). We compared the major allele frequency in the region of chromosome 18 from position 5.5 × 10^7^ onwards, finding a significantly reduced major allele frequency in clone A in both DNA (*p* = 3.7 × 10^−51^) and in RNA (*p* = 5.9 × 10^−4^), both using one-sided Wilcoxon rank-sum test.

The results of the simulations in figure 1D suggest that the higher the latent proportion of genes that exhibit CN-gene dependency, the more accurate our inference. While the set of genes that exhibit such dependency is unknown *a priori* and most likely cancer and even patient specific, we can select a set of genes that are more likely to exhibit such interactions based on previous studies. Specifically, we took the copy number and expression data from both the BRCA and OV cohorts from The Cancer Genome Atlas (TCGA, Weinstein et al. (2013)) and regressed log-expression on log *R* (relative copy number). We found the vast majority of genes exhibited a positive correlation with log *R* (supplementary figures 21 & 22). For each cancer type we selected genes with a multiple-testing adjusted *p*– value less than 0.05 as putative CN-expression interacting genes, and used these for the SA501X2B analysis though not for OV2295 as it left too few genes for stable inference.

To test the robustness of clonealign to input gene selection for the SA501, TOV2295R and OV2295R datasets we re-fitted clonealign excluding the bottom *p*% of least variable genes (as defined in log-expression space), for *p* ∈ {10, 20, 40, 60, 80, 90}, and compared the concordance in clone assignments between fits. The results can be seen as *alluvial* plots in supplementary figures 4, 12 & 13, demonstrating that clonealign is highly robust to the input gene selection and that in general up to 60% of the least variable genes may be removed before the clone assignments begin to significantly change.

To rank genes by proportion of variance explained by clonality in SA501 the full dataset was subsetted to remove any ribosomal genes and those on the X chromosome (due to entire chromosome loss). We further only considered genes whose variance in log-expression was greater than the mean variance over all genes to avoid spurious associations (ie if a gene is expressed only in a single-cell, its entire expression variation is trivially explained by clonality). The proportion of expression variation was calculated using the `aov` function in `R`. Gene Set Enrichment Analysis was then performed using the `fgsea` package (Sergushichev, 2016) using all ReactomeDB pathways with genes ranked according to proportion of expression variance explained by clonality.

## ACKNOWLEDGEMENTS

KRC is funded by postdoctoral fellowships from the Canadian Institutes of Health Research, the Canadian Statistical Sciences Institute (CANSSI), and the UBC Data Science Institute. SPS is a Susan G. Komen scholar. We acknowledge generous funding support provided by the BC Cancer Foundation. In addition, SPS receives operating funds from the Canadian Institutes for Health Research (grant FDN-143246), Terry Fox Research Institute (grants 1021 and 1061) and the Canadian Cancer Society (grant 705636). This work was supported by Cancer Research UK grant C31893/A25050 (SA and SPS).

